# Morphological limitations imposed on lizards facing urbanization

**DOI:** 10.1101/2023.05.09.540039

**Authors:** Kristopher W. Row, Oscar Villasana-Espinosa, Jaele Perez, Grace Urbina, Luke O. Frishkoff

## Abstract

Habitat conversion in general, and urbanization in particular, are thought to create ecological filters that eliminate some species while simultaneously replacing them with others that thrive under novel conditions. The specific nature of these filters is unclear, but morphology may play an important role. Here, we seek to assess which lizard morphologies are favored in urban habitats. We quantified 17 linear measurements of morphology from museum specimens from 37 lizard species from across the continental United States. We then correlate these morphological measurements with the relative incidence of observations in urban versus non-urban environments from the citizen science database iNaturalist to determine whether particular morphologies predispose species to tolerate urban environments. We then use functional diversity and morphospace volume metrics to quantify both the total amount of unique morphological diversity represented by urban associated species, versus those restricted to natural areas. Based on our results morphology appears to be filtering lizard species from urban environments. Specifically, species with intermediate body sizes and relative tails lengths, as well as larger heads and shorter hind-limbs were more likely to occupy urban zones. As a result of this filtering, there was substantially diminished morphological diversity among urban tolerant species. While natural restricted species had a high amount of morphological diversity that was unrepresented in urban tolerant species, most urban species’ morphologies were shared by natural restricted ones. Only a small subset of morphologies found in natural environments persist in urban ones, but urban lizards do possess a small number of unique morphological features that may facilitate their success. Strong selection pressures in evolutionarily novel environments are not only diminishing species diversity but pruning phenotypic diversity to favor a much smaller subset of functional possibilities. Due to the connection between phenotype and function, such diminished morphological diversity is likely to impact ecosystem functioning in impoverished human-modified systems.

## INTRODUCTION

Habitat conversion is a primary driver of biodiversity loss (Brooks *et al*., 2002; Pereira *et al*., 2012; Newbold *et al*., 2015). The environmental impacts unleashed by such conversion result in drastically altered ecological conditions that promote the loss of locally endemic species and replace them with a few expanding species that thrive in human-altered environments (McKinney & Lockwood, 1999). Urbanization is an especially drastic form of habitat modification (McDonald *et al*., 2013), and is expanding rapidly across the globe (Seto *et al*., 2013). While we have a broad understanding of how urbanization effects biodiversity and species richness, we are only beginning to understand the impact that urbanization is having on the phenotypic and morphological diversity of biological communities (Sol *et al*., 2020; Winchell *et al*., 2020).

Urban areas create massive changes in the environment, such as increases in impervious surfaces and pollution, elimination of natural vegetation, and elevated local temperatures (Shochat *et al*., 2006; Grimm *et al*., 2008). As a result of these changes, urbanization negatively affects the abundance and diversity of many native species (McKinney, 2008) while also disrupting the availability of resources that animals need to survive (Raupp *et al*., 2009). As such, the replacement of natural habitats by urban areas can precipitate large biodiversity losses—often diminishing species richness by >50% (Newbold *et al*., 2015). For example, in Southern Chile, the total number of bird species declined in urban environments due to the severe reduction in green space availability (Silva *et al*., 2016). The combination of the extreme environmental difference from natural environments and their burgeoning prevalence makes urban areas an increasingly relevant force in biodiversity declines.

However, some species buck the trend and tolerate urban areas quite well. These urban-tolerant species are likely ecologically different from urban-avoiding species along multiple niche axes. Urban species must simultaneously withstand changing food sources (Reznick & Ghalambor, 2001) altered temperatures regimes (Shochat *et al*., 2006), and broad changes in physical habitat structure (McKinney, 2002). Because morphology provides one of the key links between phenotype and fitness, many of the required shifts in niche necessary for persistence in urban environments will likely be reflected in species’ morphologies. As a result, determining whether and how morphology equates to ecological success in human-modified environments constitutes a core requirement for explaining community structure in proliferating urban ecosystems.

To date, most morphology-based research into urban-associated phenotypes has focused on quantifying the change within single species from natural to urban environments— building a case for contemporary adaptive evolution within species to novel urban habitats (Marnocha *et al*., 2011; Winchell *et al*., 2016; Putman *et al*., 2019; Putman & Tippie, 2020). For example, populations of *Anolis cristatellus* lizards in Puerto Rico possess longer limbs and more sub-digital lamallae than those in natural areas, adaptations that facilitate grasping broad surfaces such as buildings. Similarly, urban dark-eyed juncos (*Junco hyemalis),* located in southern California, have developed shorter wings and tails than their neighboring natural populations (Rasner *et al*., 2004). While evidence of such adaptation illustrates the power of strong selective pressures in individual species, the ways in which entire communities or faunas are filtered by urbanization based on their morphology remains largely unexplored (but see Sol *et al*., 2020; Winchell *et al*., 2020).

A community-wide morphological perspective has been illuminating in understanding the mechanisms behind other global change drivers: For example, in the Mediterranean, fish species with unique morphologies are most successful in invading communities (Azzurro *et al*., 2014). Elucidating community-level morphological filters determining community membership in urban areas will help contextualize species-specific results, pointing towards the generality of adaptation and pre-adaptation for survival in the city.

In this study, we assess the morphological limits imposed on lizard communities by urban environments. To do so we compare the morphological traits of lizard species that are commonly found within urban areas to lizards that rarely associate with urban zones and ask what morphologies correlate with urban success. We address three interrelated questions: First, do urban tolerant species on average come from distinct zones of morphospace in comparison to their natural counterparts? Second, do urban species take up less morphospace — indicative of only a subset of possible morphologies being viable? Finally, are morphologies that succeed in urban zones a nested subset of those in natural environments, or are urban lizards morphologically unique possessing traits that rarely occur in species restricted to natural zones? To answer these questions, we use observational data from the citizen science initiative iNaturalist, which has been shown to be a valuable resource in determining specie sensitivity to modified land use (Todd *et al*., 2016, 2017), along with the National Land Cover Database to establish the frequency with which species use urban areas. We then gather morphological data for species that span the continuum of urbanization use. By comparing morphological data to species’ frequency of occurrence in urban areas, we test the hypothesis that urbanization imposes morphological limits on lizards.

## METHODS

### Occurrence data and urban tolerance

Observational occurrence data were obtained for lizards in the continental United States from the iNaturalist database in March 2019. These data were filtered to only include research-grade observations, meaning each observation is georeferenced, has a photo, is not captive, have been reviewed and agreed upon by the iNaturalist community, had a positional uncertainty of less than 20 meters, and were observed after 2010 to ensure the observations were in line with the National Land Cover Database map used (see below). We included only species that had at least 100 post-filtering observations. The initial search yielded 67 species, of which 43 remained after filtering. Of these 43 species, on average ∼52% of the original observations remained post-filtering.

To estimate each species’ affinity to urban environments we first obtained landcover data of the United States at 30-meter resolution from the National Land Cover Database for the year 2011 (Homer *et al*., 2015). The NLCD classified urban land-covers into four categories based on the inferred amount of urban land-surface within the cell (1-19%, 20-49%, 50-79%, and 80-100% urban cover). We assigned urban values to each raster cell based on the upper level of urban cover that it contained: i.e. 80-100% received a value of 1, 50-79% received 0.8, 20-49% received 0.50, less than 20% received 0.2, and all other land-use types received a 0. We then extracted the urbanization values around the coordinates of each iNaturalist observation locality, averaging over a 100m, 500m, and 1000m radius. This buffer accommodates the 20m uncertainty tolerated around observations, and represents the degree of urbanization within the general vicinity where the observation was made and therefore the likely habitats that a lizard would encounter. The mean value for all individuals within a species constituted that species “urbanization score”, and roughly corresponds to the expected percentage of urban area within the selected radius. Comparing the averages between the different radii showed minor changes in urbanization score while not affecting the species classification as being urban or natural associated allowing the use of a 100m radius as a standard metric to determine the urbanization score of the average individual. Urbanization scores ranged from 0.0130 to 0.371, with a mean of 0.101 ± 0.084 (**Figure S1**).

Some analyses are facilitated by categorizing species into discrete groups of urban tolerant vs. non-urban tolerant. For the purpose of categorization, we applied a cutoff value of 0.1: species with urbanization scores above this cut-off value were deemed urban tolerant, and those below it were deemed natural habitat affiliated. We additionally conducted a sensitivity analysis considering alternative cut-off values (**Figure S2)**, but the overall biological conclusions are identical to those reported in the main text for a broad range of potential cut-offs.

### Morphological analysis

We assessed species’ morphology using linear morphometrics of museum specimens. Due to limitations on specimen availability, of the 43 species with urbanization scores, we collected morphology data for 37 species, and the remaining 6 species were dropped from the analysis. After identifying the five largest males for each of the 37 species (N = 185 individual lizards), we used digital calipers to make 17 individual measurements which together characterized fore- and hind-limb length, head shape, body, and tail length and shape (**Figure 1**). Not all specimens measured had complete tails, due either to (presumed) predation/competition events prior to collection, or from damage to fragile tails after preservation. Because the tail length is an important ecological feature for many lizards, we sought to estimate the total tail length of individuals with incomplete tails based on information from individuals with tails. To do so we built a linear mixed effect model to predict log total tail length as a function of log SVL, with a random effect of identity species. The conditional R^2^ of the model revealed 93% total variance of the log tail length is explained. The model allowed us to predict the total tail length of individuals with broken tails based on SVL and species identity. The predicted tail length was used as data for all individuals with broken tails (N = 23 out of 185), while real tail length was retained for individuals with complete tails.

**Figure 1:**
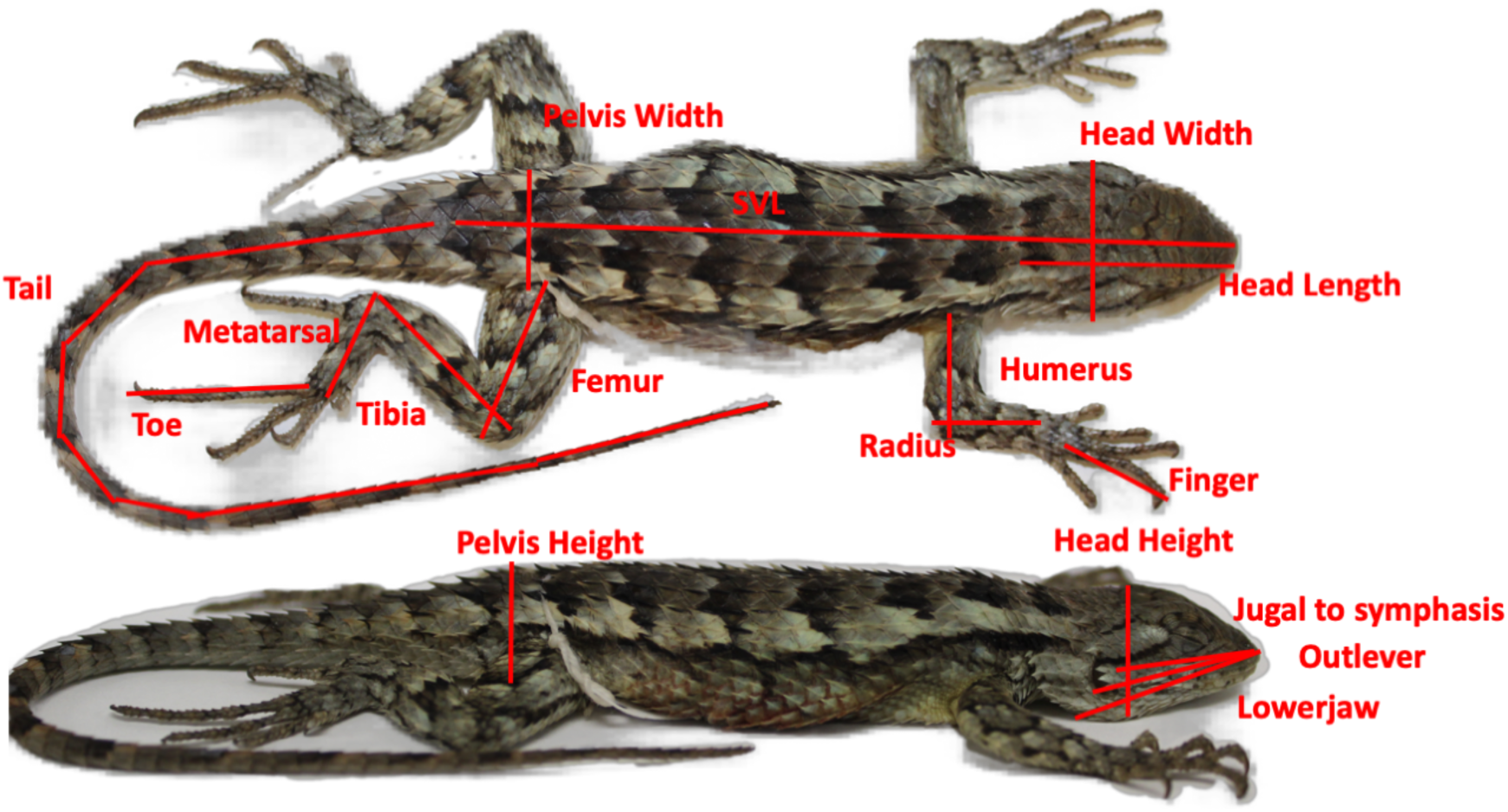
A representation of the 17 morphological traits measured, depicted against a *Sceloporus olivaceus* specimen.

To assess whether urban tolerant species occupy distinct zones of morphospace from their natural counterparts, we first ran a principal component analysis (PCA) based on the covariance matrix of all 17 morphological variables for all individuals (N = 185). All morphology measurements were log-transformed to account for body size variation being log-normally distributed. PCA reduced the dimensionality of the data by creating synthetic axes for multiple morphological traits that are highly correlated with one another. The first three principal component axes, which accounted for 95% of the total variation, were extracted and used for all subsequent analyses **(Table 1)**. To understand whether specific morphologies predispose species to tolerate urban environments we ran linear models predicting each species’ (log) urban score based on the species’ averages of the three PC axes, as well as the corresponding quadratic terms. We conducted the analysis in a multi-model framework, assessing the full model (as described previously), along with all combinations of the six predictor variables (3 linear and 3 quadratics). This multi-model framework was followed by an assessment of AIC to determine the best model. For individual models we used Wald estimates to determine the significance of model terms.

**Table 1:** Principal component axis loadings of the 17 morphological measurements across the 185 individuals and 37 species included in the study. All loadings above 0.15 and below −0.15 are color-coded to indicate the strength of positive (red) and negative (blue) correlation with each axis.

| Measurement | PC1 | PC2 | PC3 |
| --- | --- | --- | --- |
| Snout Vent Length | **0.18** | 0.04 | **0.25** |
| Head Length | **0.21** | 0.02 | **0.27** |
| Head Width | **0.24** | **-0.24** | **0.27** |
| Head Height | **0.23** | **-0.15** | **0.19** |
| Lower Jaw Length | **0.21** | 0.12 | **0.24** |
| Outlever | **0.21** | 0.10 | **0.26** |
| Jugal to Symphysis | **0.20** | 0.01 | **0.19** |
| Femur | **0.27** | -0.08 | -0.12 |
| Tibia | **0.30** | **-0.10** | **-0.33** |
| Metatarsal | **0.32** | -0.03 | **-0.45** |
| Longest Toe (4<sup>th</sup>) | **0.27** | **0.26** | **-0.40** |
| Humerus | **0.24** | **-0.24** | -0.04 |
| Radius | **0.25** | **-0.19** | -0.06 |
| Longest Finger (4<sup>th</sup>) | **0.25** | -0.01 | **-0.18** |
| Pelvis Height | **0.23** | -0.01 | **0.17** |
| Pelvis Width | **0.24** | -0.23 | **0.18** |
| Tail Length | **0.24** | **0.81** | 0.10 |
| Prop. Variation | 0.78 | 0.11 | 0.07 |

To test whether urban species take up less morphospace, we used the FD package (Legendre & Laliberté, 2010; Laliberté *et al*., 2014) to calculate two functional diversity indices: functional richness (FRic) and functional dispersion (FDis). Functional richness represents the volume resulting from the convex hull whose vertices are defined by the species in morphological space (Villéger *et al*., 2008). Functional dispersion in contrast describes how morphologically variable species in a community are, by calculating the average distance of each species from the morphological center of all species (Legendre & Laliberté, 2010).

Together, FRic and FDis provide both the total size of occupied morphospace, as well as a holistic measure of morphological variation. We use both measures because functional richness can be susceptible to outlier species. These functional diversity indices were calculated using the PC axes for both urban classified species and natural classified species (with an urban score of 0.1 applied as cutoff). To control for the differences in sample number between urban and natural species we implemented bootstrap resampling by randomly selecting between 3 and 14 species for each classification, which was repeated 100 times to create a distribution of possible functional diversity values.

Finally, we determined the level of redundancy and uniqueness between morphologies that succeed in urban zones and those in natural environments. To do so we estimated the total amount of shared morphospace between urban and natural species groups using the ‘hypervolume’ package (Blonder, 2019). This package estimates the volume and shape of the morphospace occupied and calculates the overlap and uniqueness between urban and natural morphospaces. The volume of morphospace was calculated for both urban classified species and natural classified species. The volume of unique urban morphospace was then compared to the volume of unique natural morphospace to ascertain the level at which these two species groups overlapped morphologically. Again, to control for the differences in sample size between the number of urban versus non-urban species we implemented bootstrap resampling by randomly selecting between 3 and 14 species for each classification, which was repeated 100 times to create a distribution of possible morphospace overlaps.

## RESULTS

The final dataset included a total of 185 individual specimens across 37 species. Urbanization scores ranged from 0.0130 to 0.371, with a mean of 0.101 ± 0.084 (**Figure S1**). This mean urbanization score, corresponding to the average individual of the average species occurring in an area with roughly 10% urban land cover within a 100m radius, was used as the cut-off value between classifying a species as either urban or natural. Doing so led to 14 species classified as “urban affiliated” and 23 classified as “natural affiliated”. The average body length between urban and natural lizards was broadly similar (84.2mm vs 87.5mm; t = −0.845, P = 0.34). The range of body lengths was however smaller among urban species, ranging from 50.02mm to 138.5mm, whereas natural species spanned from 49.87mm to 207.4mm.

To determine if urban tolerant species occupy distinct zones of morphospace we examined the correlation between a species’ morphology and its tolerance to urbanization. The first three principal component (PC) axes accounted for 95% of the total variation **(Figure 2a, 2b)**. The first axis (78% of morphological variation) was positively correlated with all measured morphological variables and represents overall lizard body size. Positive values of the second PC axis (11%) corresponded to species with long tails, narrow heads, and short forelimbs, while negative values indicated species with short tails, relatively wide heads, and long forelimbs.

**Figure 2:**
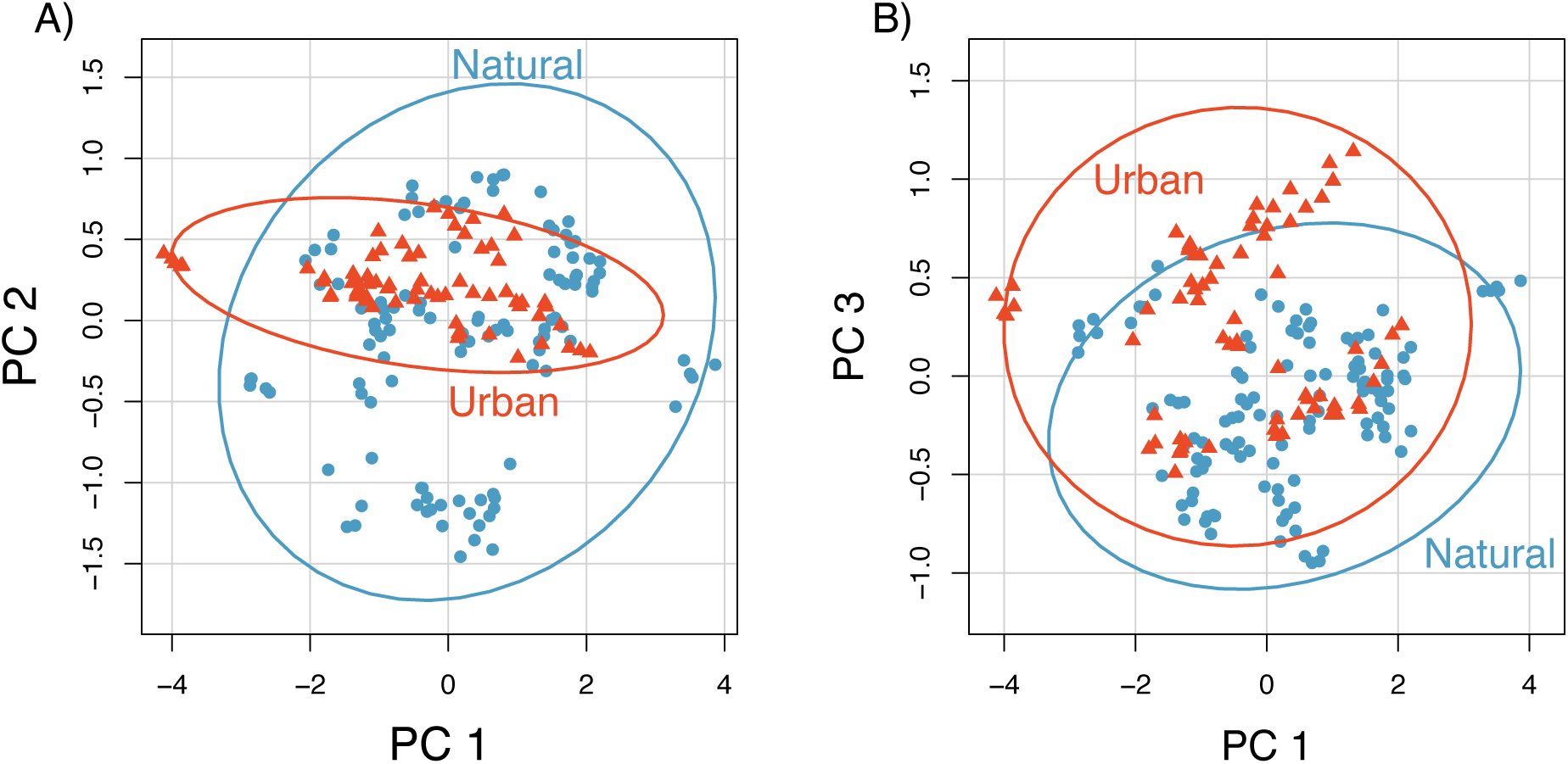
Morphological associations with urban environments. (A) Scatterplot of all individuals of all species PC1 (body size) and PC2 values (tail length, head width, fore-limb length). (B) Corresponding PC3 (body length, head size, and hind-limb length) versus PC1 (body size) values. In A and B each point represents an individual of either an urban (red) or natural (blue) affiliated species. Ellipses represent the 95% confidence intervals.

Finally, the third axis (7%) pertained to body length, head size, and hindlimb length relative to SVL, with positive values, linked to long, slender bodies, larger heads, and shorter hindlimbs. Our full model including both linear and quadratic effects of the three major morphological axes suggested that morphology strongly predicted tolerance to urbanization (F = 2.985, R2 = 0.25, P = 0.0209). Urban environments favored species of an intermediate size, with both large and small species filtered out (multiple linear regression, quadratic PC1 effect: beta=-0.083, t = − 2.111, P = 0.043; linear PC1 effect: beta = −0.145, t = −1.817, P=0.079; **Fig 3a**). Similarly, species were more likely to occur in urban areas if they had longer body lengths, larger heads and shorter hindlimbs (quadratic PC3 effect: beta = −0.360, t=−0.664, P=0.512; linear PC3 effect: beta = 0.866, t=3.129, P=0.004; **Fig 3c**).

**Figure 3:**
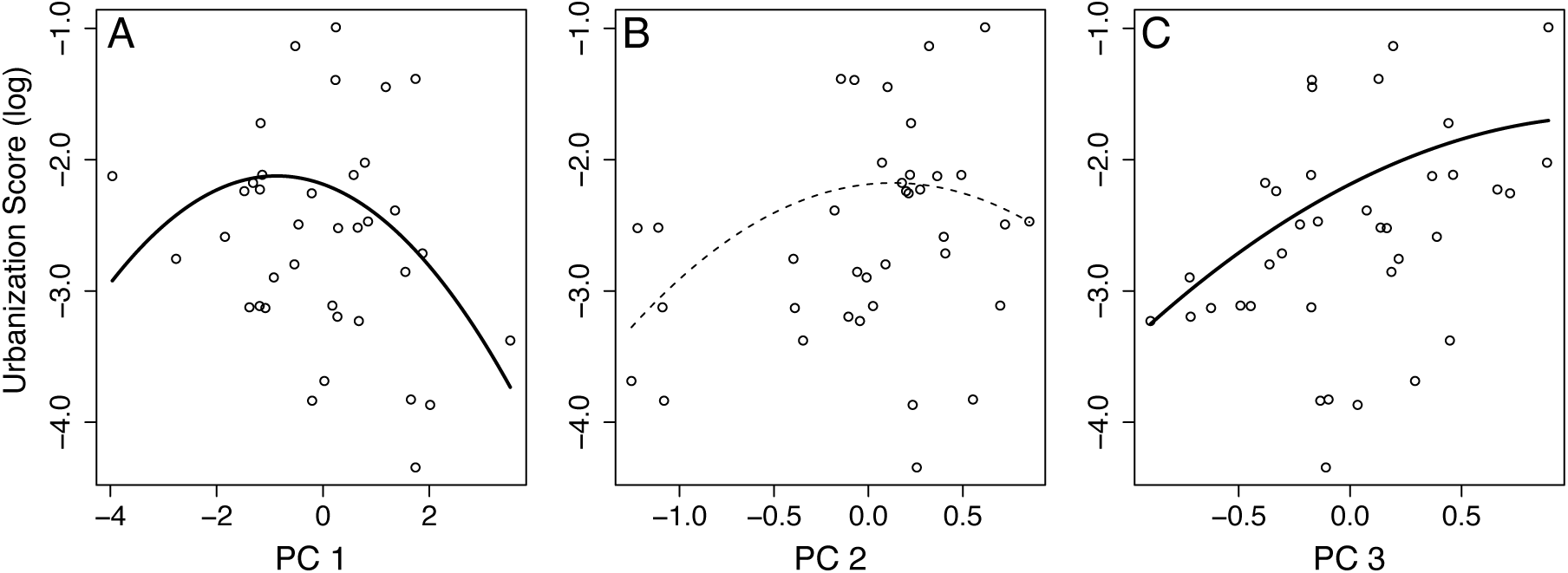
Relationships between morphology and urbanization based on the full model containing linear and quadratic effects of three major morphological PC axes. (A) Relationship between PC1 and the average amount of urban area where species is encountered (log urbanization score), depicting that intermediate-sized species are most likely to occur in urban environments. (B) Relationship between PC2 and log-transformed urbanization score. While not significant, the multimodel framework highlighted a high amount of importance in the quadratic PC2 effect. (C) Relationship between PC3 and (log) urbanization score, highlighting that urban environments favor long bodies, bigger heads, and short hind-limbs. Each point represents a species’ mean PC value. Lines depict best-fit relationship from multiple linear regression models (after taking into account the effects of other PC axes).

To verify the patterns from the full model are robust, we assessed all combinations of parameters using AIC in a multimodel framework (**Table 2**). The full model was 3.0 AIC units worse than the best-supported model. Across all models, the AIC weight of the linear component of PC3 (correlated to body length, head size, and hindlimb length) was most important (importance value: 0.92). While neither the quadratic nor the linear effect of PC2 (correlated to head size, tail length, and fore-limb length) is statistically significant in the full model (quadratic PC2 effect: beta=-0.573, t=-1.560, P=0.129; linear PC2 effect: beta=0.150, t=0.538, P=0.595), the quadratic effect of PC2 is significant in best-supported models based on AIC, and multimodel inference flags it as being relatively important (quadratic PC2 effect importance value: 0.58; **Fig 3b**). Finally, the quadratic effect of PC1 is also important (quadratic PC1 effect importance value: 0.56) which reaffirms the idea that intermediate body size is correlated to urban success.

**Table 2:** Significance and parameter values of key models evaluated in the manuscript regressing species’ (log) urbanization index score against the three major morphospace axes and their quadratic terms. ‘Rank’ displays the most likely models (all models within 2 AIC of best model) as well as the AIC ranks of the full model, and the null model against all 65 possible combinations of the six predictor variables. ‘Weight‘ indicates the AIC weight, and ΔAIC indicates the difference in AIC between the focal model and the best model. Intercept and PC terms denote parameter estimates. Significance of parameter estimates is indicated by adjacent symbols (+ p<0.1, * p<0.05, ** p<0.01). Finally, the ‘importance values’ indicate the sum of model weights over all 65 models that include each parameter.

| Rank | AIC | $\Delta AIC$ | Weight | Intercept | PC1 | PC1<sup>2</sup> | PC2 | PC2<sup>2</sup> | PC3 | PC3<sup>2</sup> |
| --- | --- | --- | --- | --- | --- | --- | --- | --- | --- | --- |
| 1 | 83.0 | 0.0 | 0.16 | -2.25 | -0.14 + | -0.08 * |  | -0.63 * | 0.84 ** |  |
| 2 | 84.4 | 1.4 | 0.08 | -2.14 | -0.15 + | -0.09 * |  | -0.70 * | 0.88 ** | -0.38 |
| 3 | 84.5 | 1.5 | 0.08 | -2.28 |  | -0.07 + |  | -0.61 * | 0.84 ** |  |
| 4 | 84.6 | 1.6 | 0.07 | -2.3 | -0.14 + | -0.08 + | 0.16 | -0.49 | 0.83 ** |  |
| 7 Full | 86.0 | 3.0 | 0.04 | -2.19 | -0.15 + | -0.08 * | 0.15 | -0.57 | 0.87 ** | -0.36 |
| 64 Null | 180.96 | 98.0 | 0.00 |  |  |  |  |  |  |  |
| Importance Values: |  |  |  |  | 0.50 | 0.56 | 0.40 | 0.58 | 0.92 | 0.21 |

To assess the volume of morphospace occupied by urban species we used functional richness (to determine the total volume of morphospace) and functional dispersion (to determine variation in morphospace). Non-urban species as a whole take up a larger amount of morphospace (FRic: 21.32) and have greater morphological variation (FDis: 1.67) than do their urban counterparts (FRic: 7.93, FDis: 1.33; **Figure 4a, 4c)**. However, because 23 species were classified as non-urban while only 14 were urban these differences could simply arise from differences in sampling depth. But even when numbers of species in each category are equalized through bootstrap resampling, the morphospace occupied by urban species is still more limited than non-urban species (**Figure 4b,4d**).

**Figure 4:**
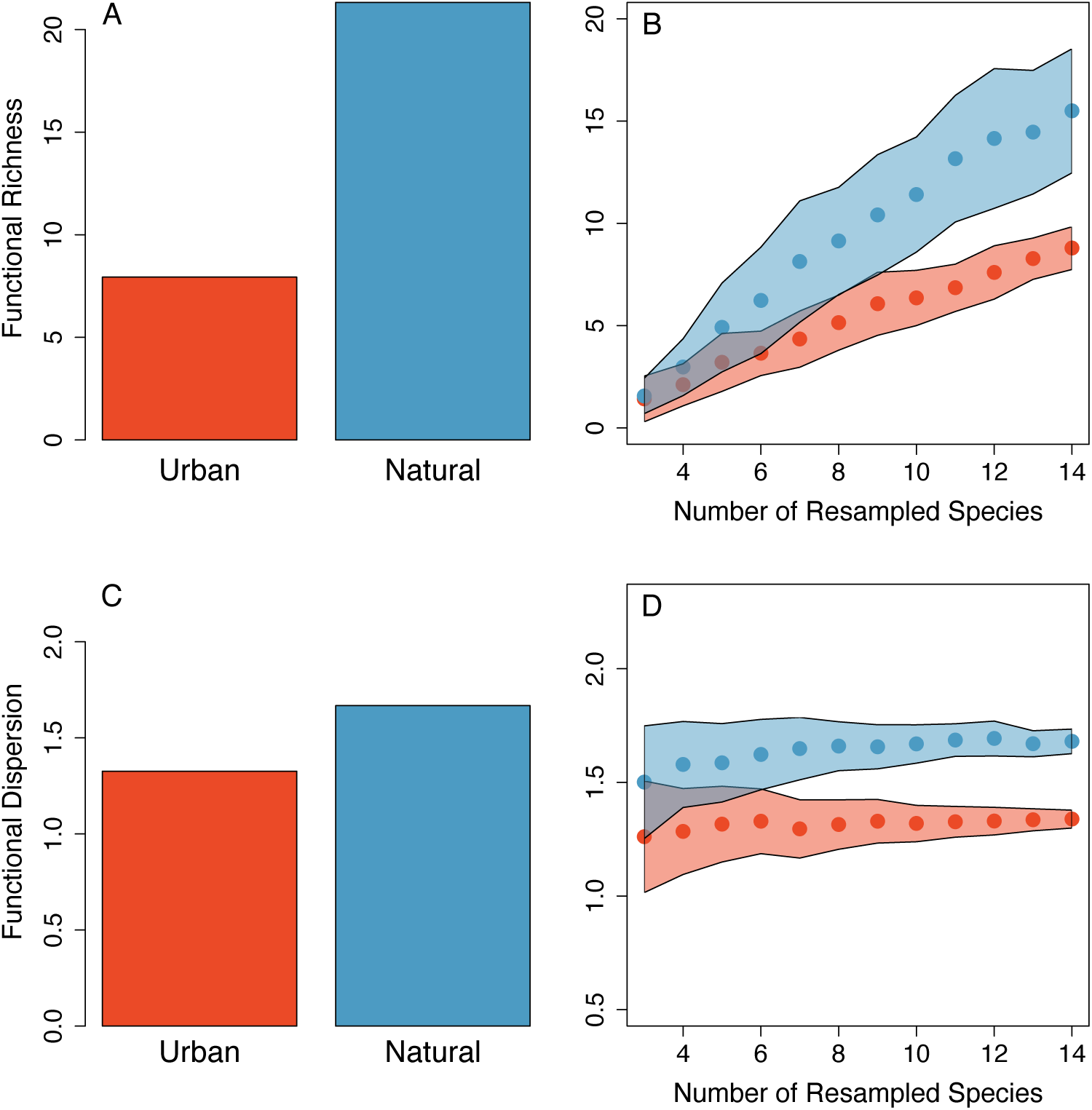
Functional diversity contained within the urban-associated and natural-restricted lizard faunas. (A) Barplots showing the total functional richness contained in urban (N = 14 species), and natural restricted (N = 23 species) faunas, with (B) bootstrap resampling used to account for a difference in sample size between faunas. Panels (C) and (D) show corresponding differences between urban and natural species for functional dispersion. In B and D points represent the bootstrapped means, and the shaded regions represent standard deviation.

Finally, to understand whether urban species contained unique morphologies, or simply represented a nested subset of morphologies contained within natural habitats, we examined the total amount of unique morphospace occupied by urban and natural species groups. When examining all species, the amount of morphospace shared by urban and natural species was relatively small (Sorensen similarity: 0.36; total overlap volume: 4.90, **Figure 5a**). Natural species possessed numerous unique morphologies—on average, 74% of morphospace occupied by ‘natural’ species did not overlap with urban morphospace (Natural unique volume: 13.97). In contrast, only 44% of morphospace occupied by urban species was unique from that of natural species (Urban unique volume: 3.83). These results were robust to resampling, such that when equal numbers of urban and natural species were analyzed, natural morphospace occupancy was 74% unique, while urban morphologies were 45% unique (**Figure 5b**).

**Figure 5:**
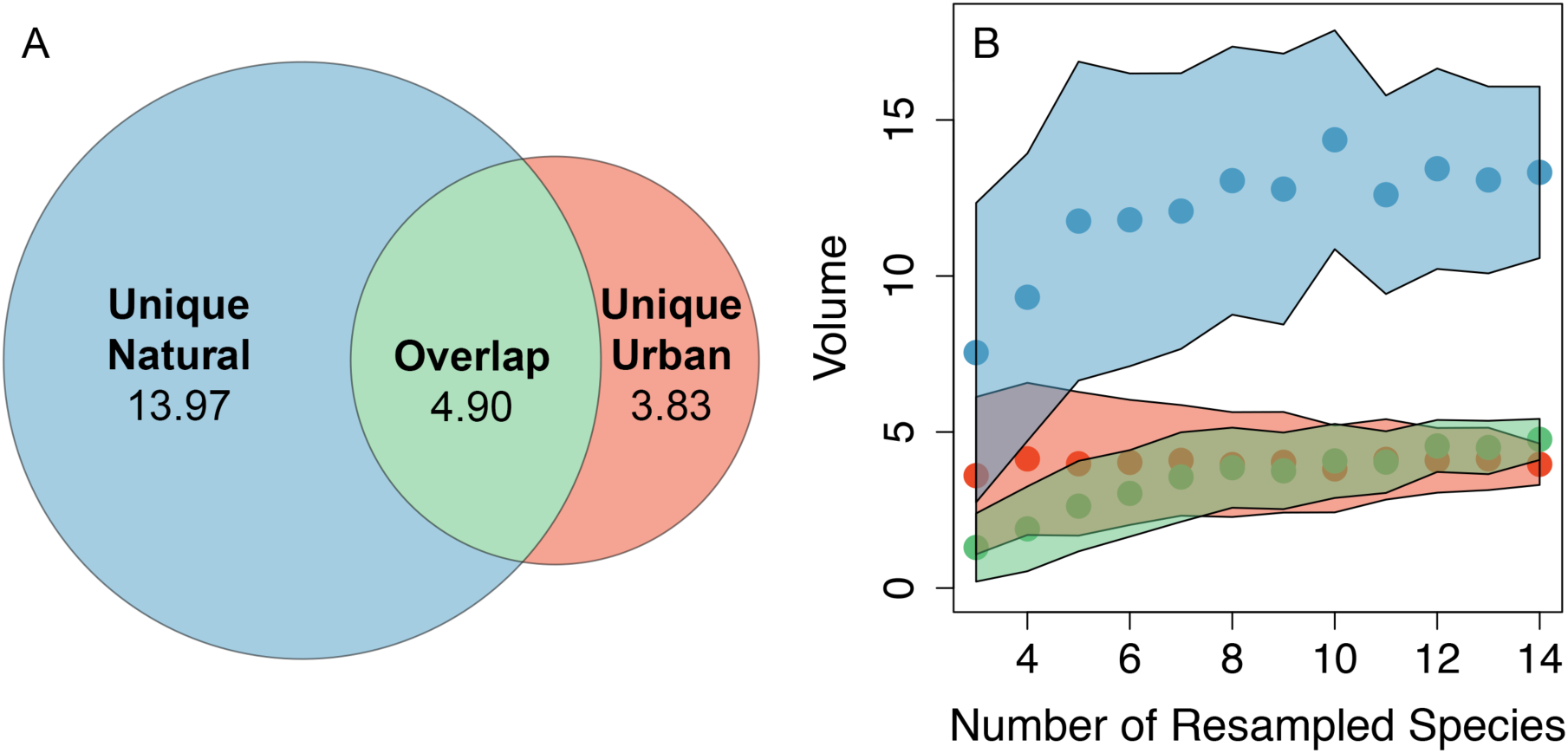
Unique and overlapping morphospace volumes occupied by urban-associated and natural-restricted lizard faunas. (A) Venn diagram showing the volume of unique and overlapping morphospace occupied by urban-associated (N = 14 species), natural-restricted (N = 23 species) faunas, with (B) bootstrap resampling used to account for a difference in sample size between faunas. In B points represent the respective the bootstrapped means, and the shaded regions around B represent standard deviations.

## DISCUSSION

We provide three key pieces of evidence that suggest that urban environments impose morphological limitations on lizard communities. First, species most prevalent in urban environments are predictable based on their morphology alone: urban areas favor species with intermediate body sizes, large heads, and short hind-limbs. Second, the morphological variation and the total amount of morphospace occupied by urban species is much smaller than that of their natural counterparts—a signal of ecological filtering, since filtering removes non-viable variation from a system. Finally, urban and natural species share only 35% of their morphospace, while >70% of morphospace occupied by natural habitat species is unrepresented among urban tolerant ones. As such, only a small subset of morphologies available in natural communities actually persist in urban environments.

While the exact reasons linking specific morphologies to tolerance of urbanization are not entirely clear, our findings suggest some likely mechanisms. Typically, large-bodied organisms are thought to be disfavored by anthropogenic impacts and are most threatened by extinction (Gaston & Blackburn, 1995; Cardillo *et al*., 2005), since body size tends to correlate with a host of life-history strategies including small clutch sizes, long times to maturity, and large home range requirements. Together these slower life-history strategies and greater requirements are thought to be poorly adapted to resource-limited environments with high potential mortality. While we do show that large-bodied lizards are absent from urban areas, their small-bodied counterparts are also excluded. The propensity for intermediate body sizes being favored could have a variety of causes. The existence of an “optimal body size” has long been postulated (Stanley, 1973; Brown *et al*., 1993), such that in the absence of competition, a given clade with a given diet and general ecology is best equipped to be a specific size due to energetic tradeoffs (Brown *et al*., 1993). A clade’s optimal body size is thought to be approximately the observed mean, as stabilizing selection pulls most species towards this mean, while competition between species for resources pushes a few species far from the mean. If species near the mean size are energetically more efficient than either small or large-bodied species, such intermediate-sized organisms may be best able to handle the stresses associated with urban life. Indeed, more recent and taxonomically expansive assessments of overall extinction risk back up the findings presented here—extinction risk is highest for both the largest and smallest species and is lower for those of intermediate size (Ripple *et al*., 2017). Other organisms experience similar reductions in body size variability in urban environments, as birds from either end of the body size distribution are also typically absent from cities (La Sorte *et al*., 2018).

Other explanations related to resource competition or physiological tolerance may however exist that specifically filter out small-bodied lizards. Thermotolerance may be essential for ectotherm survival in hotter and more variable anthropogenic environments (Frishkoff *et al*., 2015; Nowakowski *et al*., 2018b), but small-sized lizards retain less heat, directly affecting their ability to thermoregulate (Michael *et al*., 2014). Alternatively, resource limitation and both interference and exploitative competition may play a role in explaining the failure of smaller species. Most lizards are generalist insectivores and select prey based on their size. Insect abundance is severely reduced in urban environments (Merckx *et al*., 2018). Due to gape limitation, intermediate-bodied species have access to a larger range of prey sizes (both small prey and large prey), while smaller lizards are forced to compete for small prey both amongst themselves and with larger lizards (Herrel *et al*., 1995; Lima *et al*., 2000; Vitt, 2000). This mechanism would also explain why species with large head sizes are preferentially abundant in urban areas. Interference competition between species in urban environments may similarly play a role, as larger heads are also paramount in aggressive displays, both within and between species (Donihue *et al*., 2016; Wegener *et al*., 2019). More behaviorally aggressive species and individuals are often more common in human-modified environments, with the best cases coming from studies of birds (Shochat *et al*., 2010; Scales *et al*., 2011; Hernández-Brito *et al*., 2014). For example in agricultural landscapes, birds feedings at isolated trees are primarily aggressive dominant species with large bill sizes, while subordinate species are restricted to trees near forest-agriculture ecotones (Daily & Ehrlich, 1994).

Limb length reflects a trade-off between agility and speed on broad surfaces versus narrow surfaces. In general, longer hind-limbs are associated with faster-running speeds and ecologies in which an individual needs to flee from predators and run-down prey. Shorter limbs in contrast grant individuals the ability to navigate narrow and irregular surfaces, greater climbing ability, and are often associated with sit-and-wait predator strategies (Losos, 2009). Urban environments are dominated by buildings and older trees, and possess a denuded understory—features that in some ways mimic natural environments where the need to climb benefits lizards with shorter limbs (Herrel *et al*., 2001). This trend was also directly observed in western fence lizards where females in urbanized environments had shorter limbs than their non-urban counterparts (Sparkman *et al*., 2018). However, urban populations of *Anolis cristatellus* possess longer limbs than natural populations (Winchell *et al*., 2016), indicating that the benefits of short limbs are not ubiquitous and that multiple eco-morphological strategies might exist to maximize fitness in urban environments. Alternatively, the preference for short limbs may come about due to its association with sit-and-wait foraging, a strategy that in contrast to active foraging, may avoid risks from enhanced predation from mesopredators, like feral cats, or pseudo-predation from automobiles. Indeed our findings are consistent with some emerging patterns of urban success found in other lizard faunas. In the Caribbean urban tolerance is negatively associated with relative hind limb length across the genus *Anolis* (Winchell *et al*., 2020)

At the level of the “urban lizard fauna”, we find that the total morphological diversity represented in these urban systems is much smaller than that of the primarily natural fauna. Such a reduction of morphological variation among the urban assemblages mirrors general losses of functional diversity (of which morphological diversity is a subset) that are frequently observed in human-modified environments. For example, urban bird assemblages show an average decrease of 20% in functional diversity when compared to their surrounding natural habitats (Sol *et al*., 2020), and for birds and mammals, functional diversity declines sharply as agricultural land use intensifies (Flynn *et al*., 2009). Among these North American lizards, the reduction in morphological variability is partially attributable to large and small-bodied lizards being preferentially excluded from urban areas. While overall the urban tolerant species group contained less overall morphological diversity, the remaining diversity was not a simple subset of the diversity among the natural habitat affiliated species. Instead, roughly 40% of the morphological diversity of urban species was unique. This finding does not necessarily mean that there are truly unique morphotypes in urban environments, as even the most urban affiliated species in this dataset still occur frequently in natural environments as well. Addressing whether urban environments typically contain morphologically unique species that are absent or rare in nearby natural environments would require standardized surveys of individual lizard communities, a task that presence-only citizen science occurrence data is not suited for. Regardless, the overall reduction in morphospace occupation, along with the shift towards some unique morphologies among the most urban tolerant lizards, suggest that urban environments represent a strong selection pressure at the community level.

The ability of anthropogenic pressures to reduce diversity beyond the taxonomic level has become more widely recognized as a conservation challenge (Devictor *et al*., 2010). In particular, phylogenetic and functional diversity a multiple scales are eliminated by intense forms of anthropogenic change (Sol *et al*., 2020). often above and beyond that expected from species loss alone (Flynn et al., 2009; Frishkoff et al., 2014; Hagen et al., 2017). Such diversity declines are especially worrisome because functional diversity is more tightly linked with ecological functioning (Tilman *et al*., 1997), and its decline therefore likely portends loss of the services that ecosystems provide to humans (Karp *et al*., 2013; Monagan *et al*., 2017; Echeverri *et al*., 2021). Whether reductions in lizard diversity observed here result in meaningful declines in the services lizards provide, such as pest control, remain unknown. However, lizard abundance has been linked with agricultural insect pest consumption (Monagan *et al*., 2017), suggesting that their role as service providers may be relevant in urban areas.

Together our data show that the urban lizard fauna is a morphologically restricted set of species, implicating the urban environment as imposing ecological filters on urban community assembly. Importantly we go beyond simply demonstrating morphological diversity losses, but show specifically how changes in morphological composition results in this diversity loss— namely through reduction of large and small body sizes and elimination of lizards with smaller heads and longer limbs. While our analysis shows that morphology predicts roughly 25% of the variation in relative affiliation with urban versus natural environments, other unconsidered features likely also play a role. For example, species with higher trophic positions, or those that depend on aquatic habitats may be especially vulnerable to changing land-use (Todd *et al*., 2016, 2017). Critically however, many of these ecologies will be reflected in species morphologies. However, other traits will not be reflected in morphology. Of these, physiological traits, especially heat tolerance, are likely to be paramount, as thermal tolerance has been repeatedly linked with success in human-modified systems (Frishkoff *et al*., 2015; Nowakowski *et al*., 2018a).

This study highlights an analytical framework to assess the prevalence of morphological filters in human-dominated landscapes using widely available, and ever-increasing citizen science data. Whether the patterns documented here are general features of urban systems across taxa or across geographic space remains to be seen. The traits that predict tolerance to other forms of anthropogenic change are sometimes inconsistent between regions (Hatfield *et al*., 2018) or types of habitat conversion (Bartomeus *et al*., 2018). Indeed the same species traits may increase tolerance to land-use change in some climate zones while decreasing tolerance in others (Murray *et al*., 2021). As more of the filters determining community composition in anthropogenic systems are understood, the ecological rules that define the prevailing biological communities of the Anthropocene will come into focus. Hopefully, with an appreciation of these rules, targeted interventions to make anthropogenic systems more wildlife-friendly can be implemented, which will support species with traits that were previously filtered out.

## Supporting information

Raw_data_spreadsheet

## Acknowledgements

We would like to thank the museums and biodiversity collections that made collecting the lizard morphological data possible. Specifically, we thank Carol Spencer, Michelle Koo, as well as the staff of the UC Berkeley Museum of Vertebrate Zoology for allowing access to the herpetology collection. Additionally, we thank the faculty and staff that support the Amphibian and Reptile Diversity Research Center at the University of Texas at Arlington for their instructions on proper museum etiquette and procedures.

## Supplemental Figures

**Figure S1:**
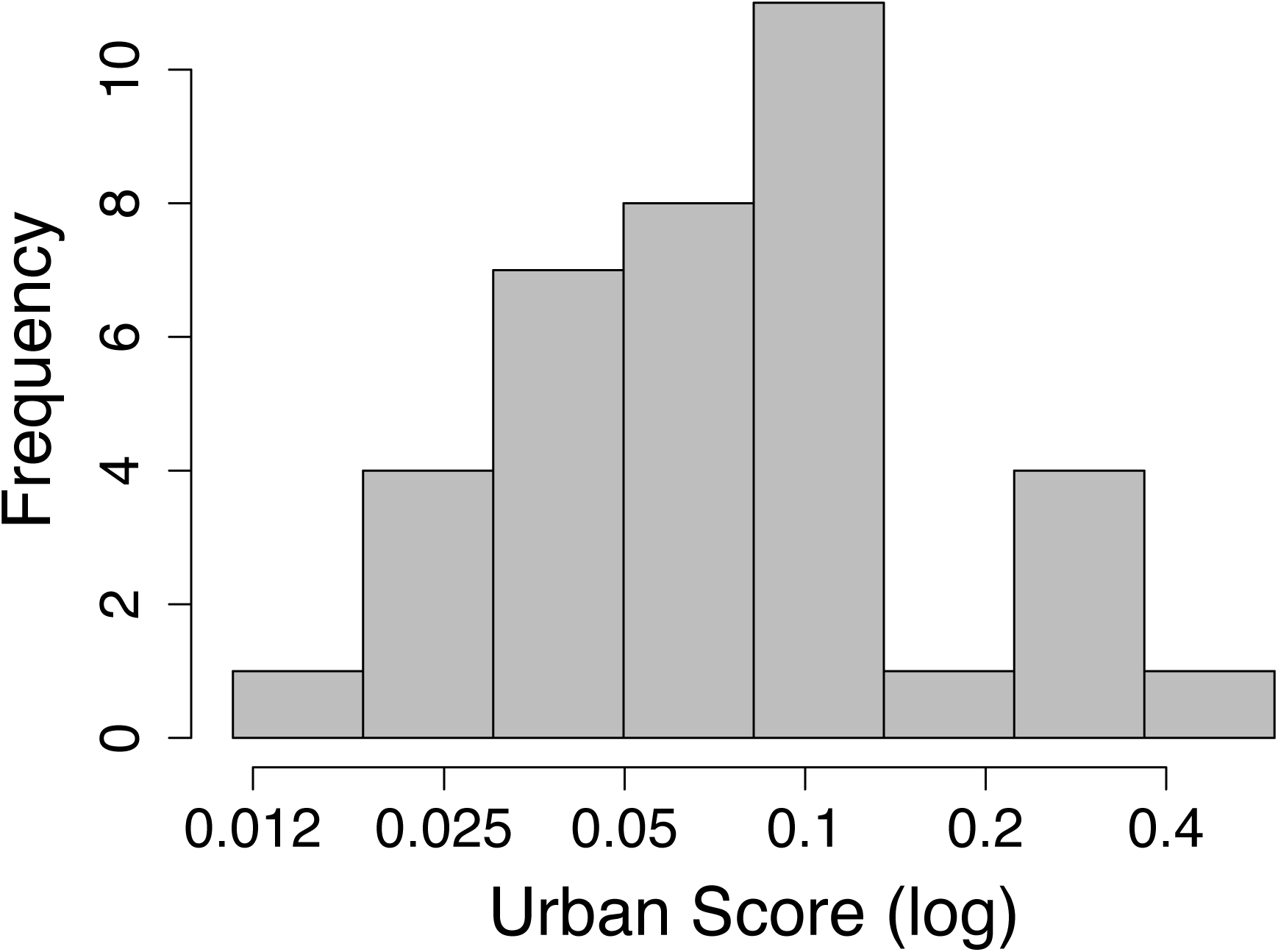
Histogram showing the distribution of urbanization score across all 37 species used in the analysis. Note that x-axis is on log-scale.

**Figure S2:**
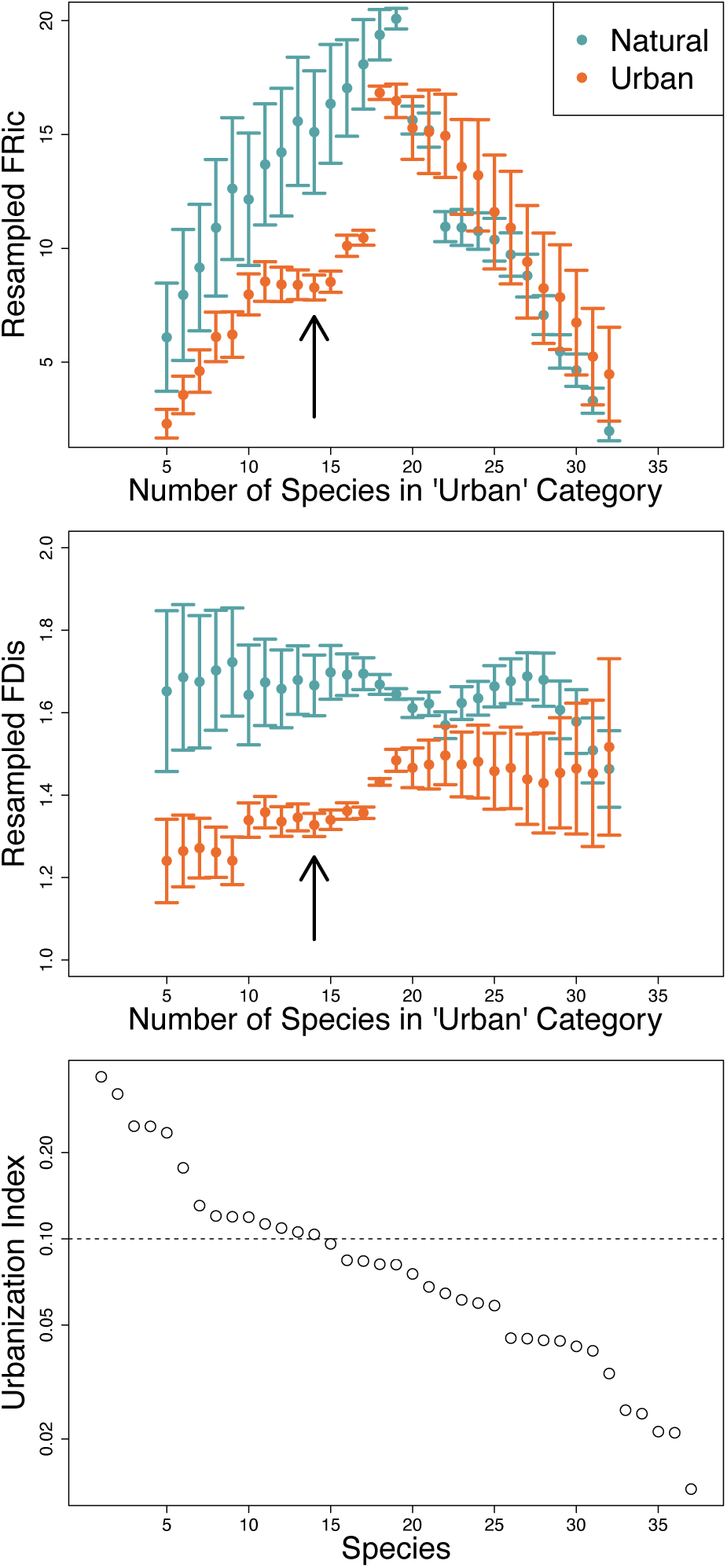
Sensitivity analysis evaluating the consequences of alternative cut-off values to determine “urban” species. In the top two panels functional richness (FRic) and functional dispersion (FDis) are evaluated under alternative cut-off values by considering the X most “urban” species, based on their urbanization index value, and then resampling (without replacement) the remaining (more natural affiliated) species down to X, and calculating Fric and FDis on this equal-sized subset. We evaluated all values of urbanization cut-offs, starting with the five most urbanized species up to 32 when there were only 5 species in the “natural” category. In top and middle panel points depict means and lines show the standard deviations over 1000 resamplings for each value of X. Black arrow shows the number of species considered “urban” in the main text. The lower panel shows the corresponding urbanization values for all species in the dataset, ordered from most to least urban affiliated, such that placement along the x-axis corresponds to the resampled values of species in the upper two panels. The dashed line shows the urbanization cut-off (0.1) used in the main text. Note that the y-axis is logarithmic.

